## Supplementary Figures for "SLC25A46 localizes to sites of mitochondrial fission and fusion and loss of function variants alter the oligomerization states of MFN2 and OPA1"

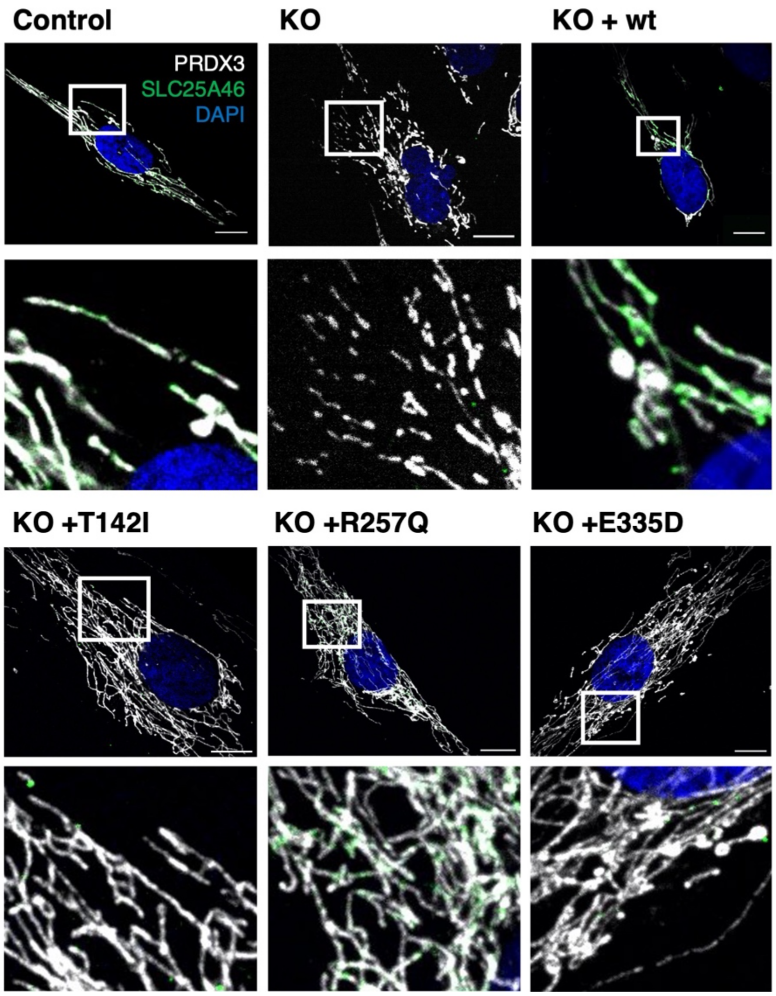


**Supplementary Figure 1: Expression of the SLC25A46 pathogenic variants.**

Control fibroblasts, the knock-out cell line, cells with reintroduced SLC25A46 protein (WT, p.T142I, p.R257Q and p.E335D) were analyzed by immunofluorescence. Representative images of fibroblasts decorated with anti-PRDX3 (mitochondria) in white, DAPI (nuclei) in blue and SLC25A46 in green. Scale bars: 10 $\mu$m

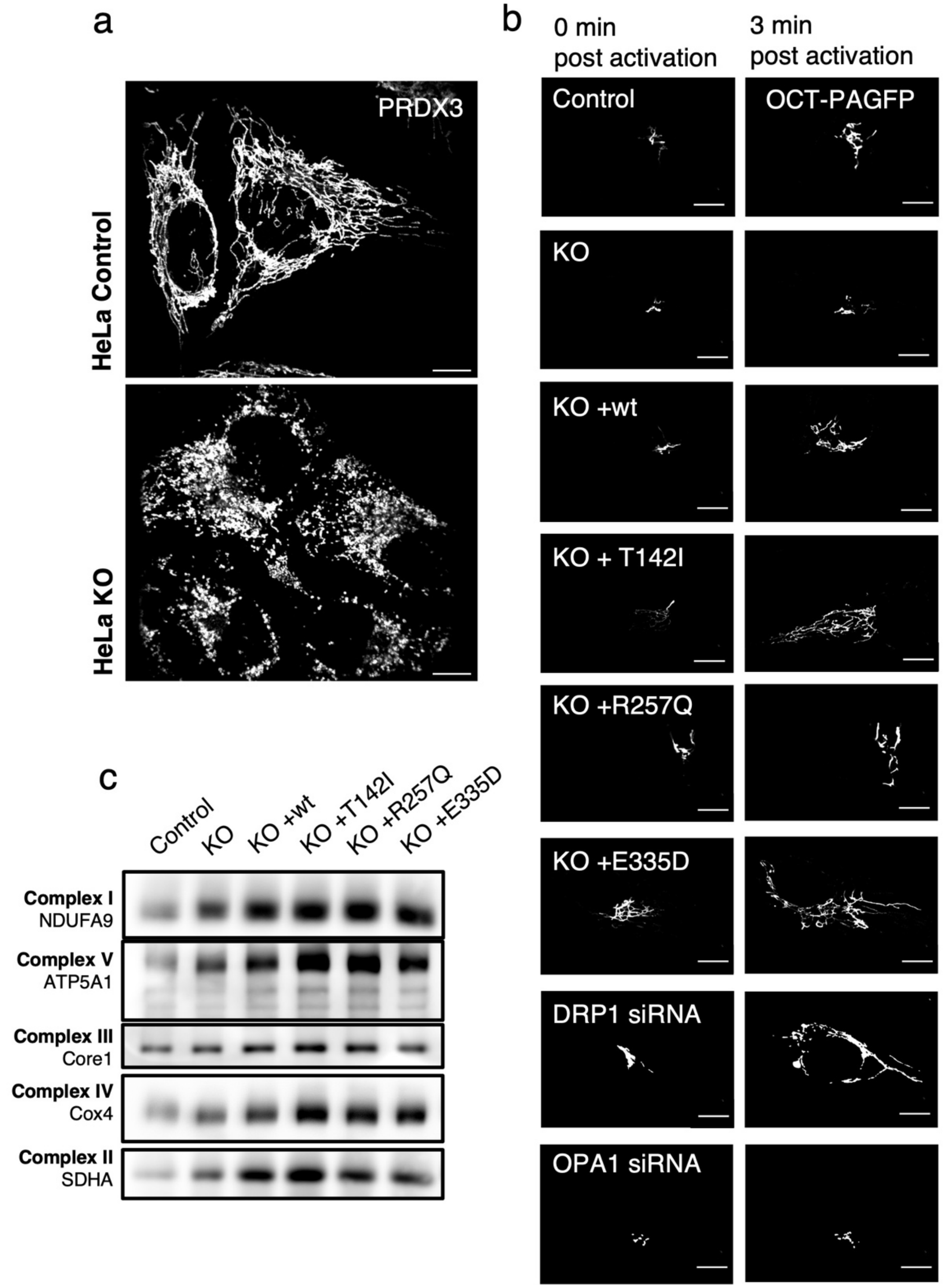


**Supplementary Figure 2: Physiological consequences of SLC25A46 knock-out and expression of pathogenic variants.**

a) The mitochondrial morphology of HeLa control cells and knock-out of SLC25A46. Representative images of cells decorated with anti-PRDX3 (mitochondria) in white. Scale bars: 10 $\mu$m. b) Live-cell imaging analysis of fibroblasts expressing a mitochondrial targeted photoactivatable GFP (OCT-PAGFP) probe. Representative images of the OCT-PAGFP probe diffusion over a 3-min period after activation with the 405 nm laser. Scale bars: 10 $\mu$m. c) BN-

PAGE analyses of OXPHOS complexes in fibroblasts from control and SLC25A46 knock-out cell line, and cells expressing the wildtype protein or the pathogenic variants of SLC25A46. Each of the five OXPHOS complexes (I–V) was visualized with a subunit-specific antibody that recognizes the native complex.

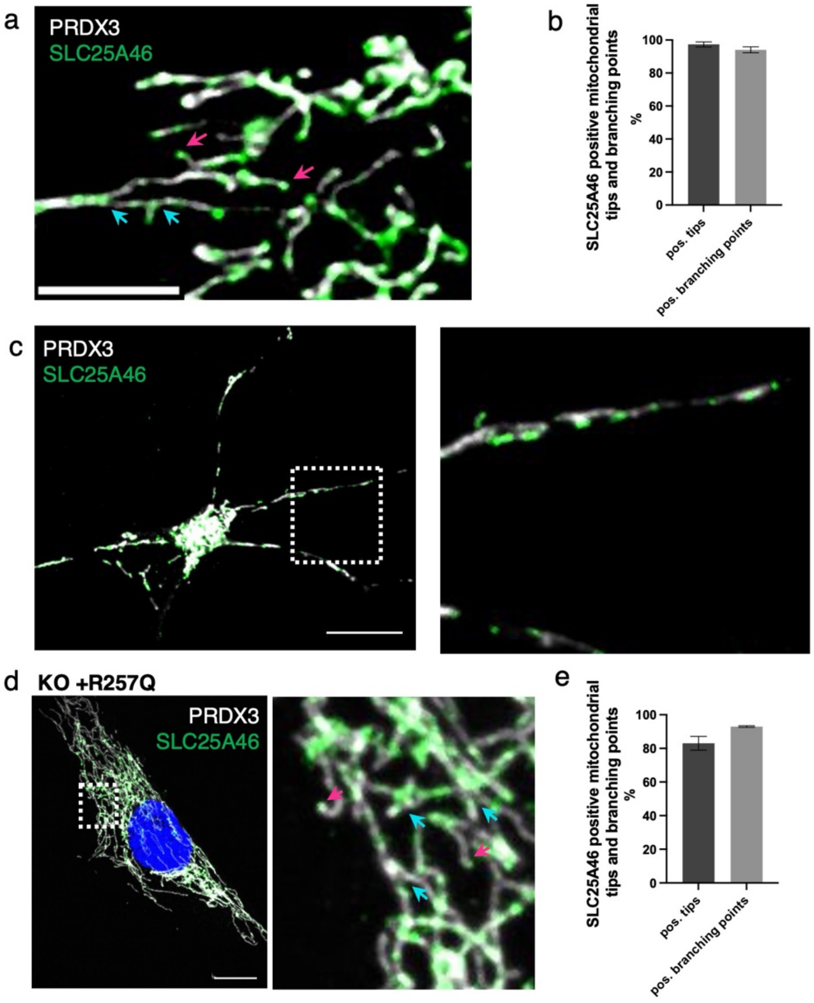


**Supplementary Figure 3: Endogenous SLC25A46 is present at mitochondrial fusion and fission sites.** (a) Immunofluorescence analysis of human fibroblasts. Endogenous SLC25A46 localization is shown in green. PRDX3 is used as a mitochondrial marker (white). Pink arrows indicate mitochondrial tips, blue arrows indicate mitochondrial branching points. Scale bar: 5 μm. (b) Mitochondrial tips (n>80 per condition) and branching points (n>33 per condition) were analyzed for positive focal SLC25A46 signal. Data represents mean+SEM, n=3. (c) Immunofluorescence analysis of cortical neurons differentiated from human iPSCs. Endogenous SLC25A46 localization is shown in green. PRDX3 is used as a mitochondrial marker (white). A zoomed image of the box area is shown on the right. Scale bar: 10 μm. d) Immunofluorescence analysis of SLC25A46 knock-out fibroblast cell lines expressing the R257Q variant. SLC25A46 localization is shown in green. PRDX3 is used as a mitochondrial marker (white). Pink arrows indicate mitochondrial tips, blue arrows indicate mitochondrial branching points. A zoomed image is shown on the right. Scale bar: 10 μm. e) Mitochondrial tips (n>60 per condition) and branching points (n>30 per condition) were analyzed for positive focal SLC25A46-GFP signal in fibroblasts expressing the pR257Q variant. Data represents mean+SEM, n=3.

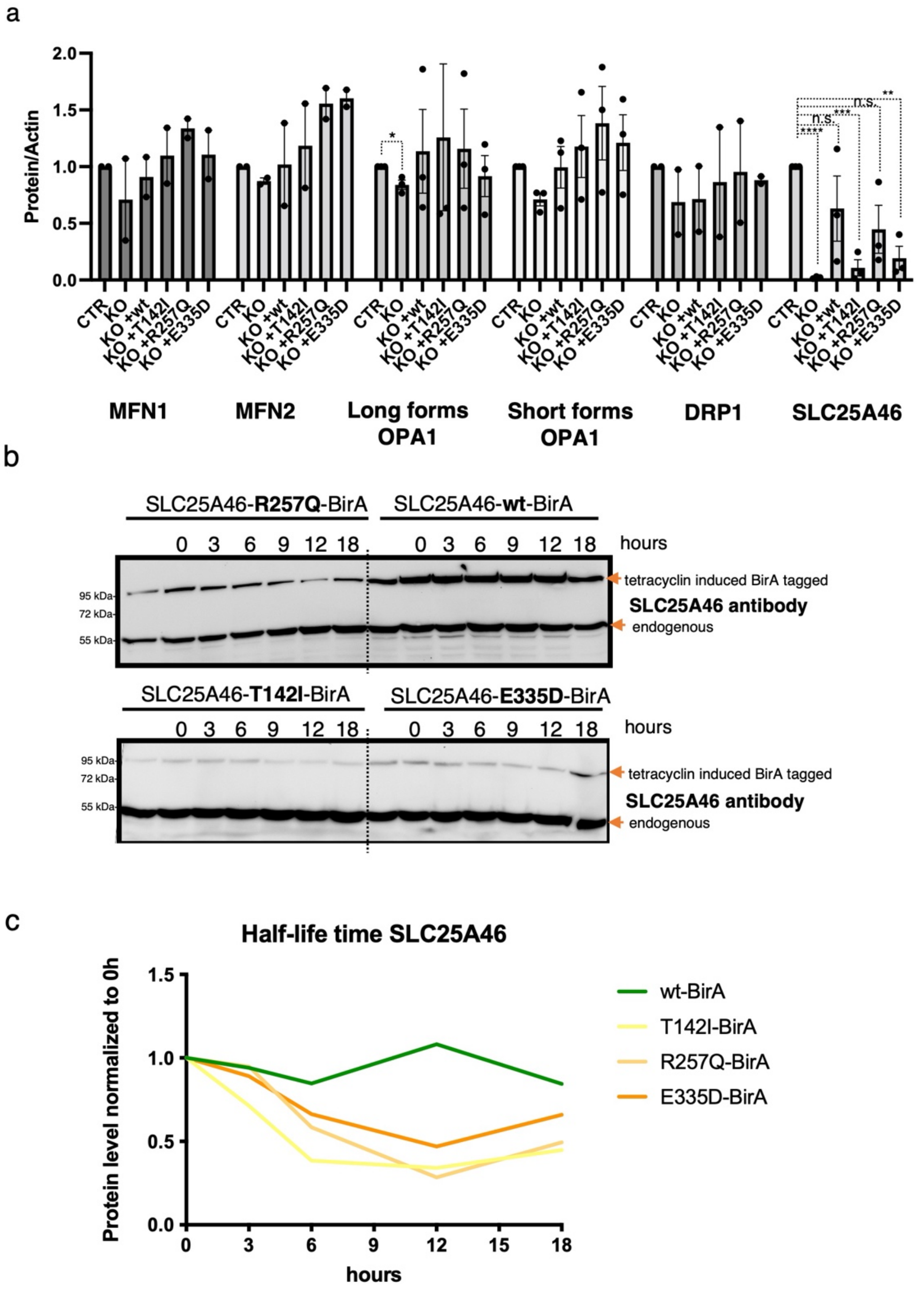


**Supplementary Figure 4. No difference in steady-state protein levels of fusion and fission proteins and decreased half-life of SLC25A46 pathogenic variants**

a) Quantification of SDS-PAGE analysis (as shown in Fig. 5A) of control fibroblasts, SLC25A46 knock-out and knock-out overexpressing wildtype SLC25A46 or pathogenic variants (T142I, R257Q, E335D). Actin serves as a loading control. P-values were calculated using a two-tailed, unpaired t-test *P<0.05, n=3 independent experiments. If not indicated, no significant differences were found. b) Flp-In T-REx 293 cells expressing inducible SLC25A46 (wt) or the pathogenic variants (T142I, R257Q, E335D) with a BirA* tag were incubated with tetracyclin for 24 h and washed thoroughly. The cells were harvested after indicated hours and protein expression of BirA*-tagged proteins was measured by Western blotting using SLC25A46 antibody. c) Quantification of SDS-PAGE in b) normalized to endogenous expression of SLC25A46 and 0 h.

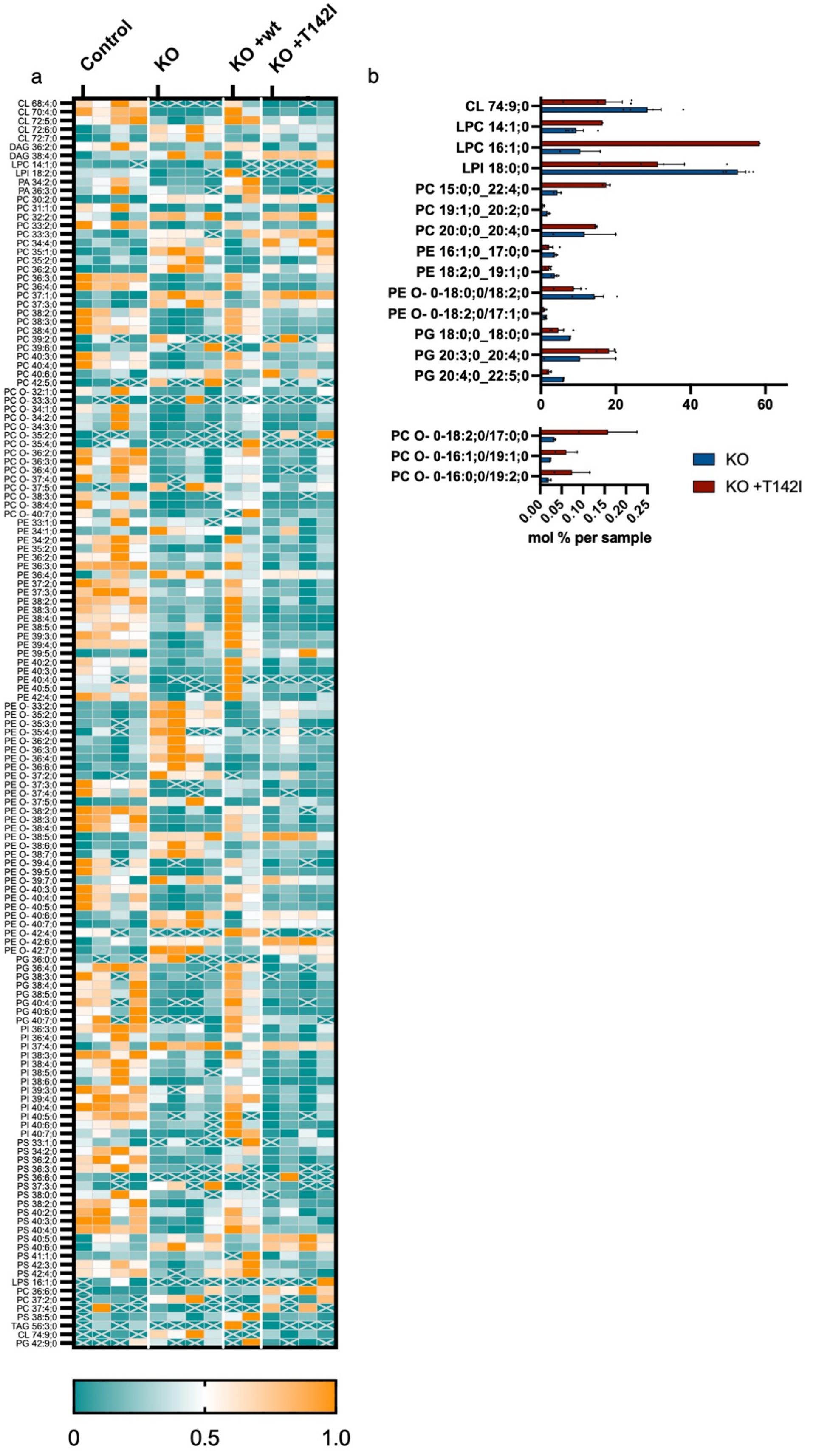
a
Control
KO
KO +wt
KO +T142I
0
0.5
1.0
b
CL 74:9;0
LPC 14:1;0
LPC 16:1;0
LPI 18:0;0
PC 15:0;0_22:4;0
PC 19:1;0_20:2;0
PC 20:0;0_20:4;0
PE 16:1;0_17:0;0
PE 18:2;0_19:1;0
PE O- 0-18:0;0/18:2;0
PE O- 0-18:2;0/17:1;0
PG 18:0;0_18:0;0
PG 20:3;0_20:4;0
PG 20:4;0_22:5;0
0
20
40
60
PC O- 0-18:2;0/17:0;0
PC O- 0-16:1;0/19:1;0
PC O- 0-16:0;0/19:2;0
0.00
0.05
0.10
0.15
0.20
0.25
mol % per sample
KO
KO +T142I

**Supplementary Figure 5: Loss of SLC25A46 leads to alterations in mitochondrial lipid subspecies**

Sucrose bilayer purified mitochondria from control human fibroblasts (control, n=4), SLC25A46 knock-out fibroblasts (KO, n=4) and SLC25A46 knock-out fibroblasts expressing either the wildtype protein (KO +wt, n=2) or the pathogenic variant T142I (KO +T142I, n=4) were analyzed for absolute quantification of lipid content using shotgun mass spectrometry lipidomics. a) Heat map representing relative abundance of 203 lipid entities that were significantly different. Data were normalized within each lipid species. b) Subspecies of the significantly different lipids found between SLC25A46 knock-out fibroblasts (KO) and SLC25A46 knock-out fibroblasts expressing the pathogenic variant T142I (KO +T142I). Values are indicated as mol % of sample.

**Supplementary Video 1: SLC25A46 is present at fusion and fission sites**

Videos of the captures in Fig. 3. Fibroblasts stably over-expressing SLC25A46-GFP (green) were analyzed in live-cell imaging. Mitochondria were stained with MitoTracker Deep Red (white). Images were captured every 0.5 seconds for a period of 1 minute. (fusion) A magnification of the time-lapse imaging of a fusion event. (fission) A magnification of the time-lapse imaging of a fission event.
